# The rate of whole-genome duplication can be accelerated by hybridization

**DOI:** 10.1101/2020.10.14.339820

**Authors:** S. Marsit, M. Hénault, G. Charron, A. Fijarczyk, C. R Landry

## Abstract

Hybridization and polyploidization are powerful mechanisms of speciation. Hybrid speciation often coincides with whole-genome duplication (WGD) in eukaryotes. This suggests that WGD allows hybrids to thrive by restoring fertility and/or increasing access to adaptive mutations. Alternatively, it has been suggested that hybridization itself may trigger WGD. Testing these models requires quantifying the rate of WGD in hybrids without the confounding effect of natural selection. By measuring the spontaneous rate of WGD of 1304 yeast crosses evolved under relaxed selection, we show that some genotypes are more prone to WGD and WGD can be triggered by hybridization. We also find that higher WGD rate correlates with higher genomic instability and that WGD increases fertility and genetic variability. These results provide evidence that hybridization itself can promote WGD, which in turn facilitates the evolution of hybrids.

## Main text

Whole genome duplication (WGD) is an important evolutionary force in eukaryotes ^1–6^. While significant progress has been made in the last decade in understanding the consequences of WGD ^2,7^, the factors that favor its evolution are less well studied ^8^. A large body of work, mostly on plants, showed that hybrid speciation often coincides with WGD ^2,9,10^. A long-standing hypothesis suggests that parental genetic divergence influences the probability of WGD in hybrids ^11–13^. Combining two diverged genomes in the same cells would increase the rate of genomic changes and WGD, which would in turn enable hybrid maintenance on the long term ^13^. However, the role of genetic divergence as a driver for WGD has since been debated ^14–19^. The debate comes from the fact that hybrid species could be more likely to be maintained because WGD increases fitness by restoring fertility ^11,20–22^ and accelerating adaptation ^23^, rather than hybridization itself increasing the rate of WGD. Testing these alternative models requires to remove natural selection from the equation. Here we show, using yeast as an experimental model, that WGD is more likely to occur in some lineages, but can also be triggered by hybridization through the combination of some genotypes. We also find that higher WGD rate correlates with higher genomic instability, and that WGD increases fertility and genetic variability. Together, these results provide evidence that hybridization itself can promote WGD.

Using yeast as an experimental model, we investigated whether hybridization can trigger whole-genome duplication (WGD) under relaxed selection. We measured the rate of WGD in 1304 independent yeast lines from 15 crosses following a protocol for mutation accumulation (MA). MA lines were evolved through repeated strong bottlenecks to remove the confounding factor of natural selection. We used crosses over different levels of parental divergence from intra-lineage to inter-specific crosses (Fig. 1A). Natural yeast isolates representing three incipient species of the wild species *Saccharomyces paradoxus* and its sister species, *S. cerevisiae*, were used. The *S. paradoxus* lineages (*SpA, SpB* and *SpC*) exhibit up to 4% nucleotide divergence, and 15% with *S. cerevisiae* while their genomes remain largely co-linear ^24,25^ (Fig. 1A). We previously generated 864 lines ^22,26^ and here generated 288 new lines (3 crosses) using the same procedure (Table S1). All crosses were evolved for roughly 770 generations (Fig. 1A). We also examined 152 MA lines of a diploidized haploid *S. cerevisiae* strain propagated for ∼2,062 generations ^27,28^. The MA lines were classified in terms of nucleotide divergence in four types: Very Low (VL: VL_B = *SpB* × *SpB*, VL_C = *SpC* × *SpC*, VL_A = *SpA* × *SpA* and VL_S = *Scer* × *Scer*); Low (L = *SpB* × *SpC*); Moderate (M = *SpB* × *SpA*) and High (H = *SpB* × *S. cerevisiae*) (Fig. 1A, Table S1).

**Fig. 1.**
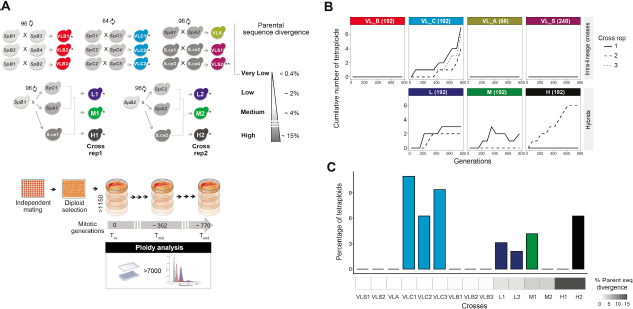
Whole genome duplication is genotype specific and can be accelerated by hybridization. (A) Crosses among *S. paradoxus* incipient species (VL_B, VL_C, VL_A, L, and M), among *S. cerevisiae* isolates (VL_S) and between these two distant species (H). Most types of crosses involve two to three biological replicates, and each individual cross was performed independently 48-96 times to represent independent hybridization. Crosses are labeled with a star according to their source *(Charron et al., 2019; Hénault et al., 2020) and **(Hall et al., 2008). The 1304 lines were evolved under relaxed selection. Mitotic propagation was performed through repeated single cell bottlenecks and ploidy was measured using flow cytometry at different generation timepoints. (B) Cumulative number of tetraploid lines observed along the experiment. Numbers in parentheses represent sample sizes. (C) Whole genome duplication rate differs among crosses (VL_C1 (n = 7), VL_C2 (n = 4), VL_C3 (n = 6), L1 (n= 3), L2 (n= 2), M1 (n= 4), H2 (n= 6)).

We measured the change in ploidy of all lines by quantifying DNA content using flow cytometry. We detected WGD (from diploid to tetraploid) in 7 crosses of all 4 types (Fig. 1B, fig. S1). Thirty-two tetraploid lines were identified in total (fig. S2). Using whole genome sequencing, we found that, for all tetraploids, the frequency of parental alleles across the genomes is roughly 50%, confirming that both parental genomes have been entirely duplicated (fig. S3). This shows that within 770 generations, 0 to 11% of the lines went through spontaneous WGD (Fig. 1B, C, Table S2).

WGD occurred in the 3 hybrid types (L, M and H) and in only one of the intra-lineage crosses (VL_C). Since WGD is observed exclusively in the three VL_C crosses (VL_C1 (11%), VL_C2 (6%), VL_C3 (9%)) and not the other VL crosses, we set out to investigate if *SpC* strains are intrinsically more prone to WGD. To test this, we analyzed the ploidy of more than 300 wild North American *S. paradoxus* isolates belonging to the three *SpA, SpB* and *SpC* lineages ^24^. Consistent with the observations in the MA lines, *SpC* is the only lineage from which a natural tetraploid strain was found (fig. S4). We have also previously shown that some *SpC* haploid strains are prone to spontaneous diploidization (1n to 2n) ^22^. As a consequence, almost half of the L1 and L2 lines are triploid (fig. S1) that most likely arose from mating between pseudo-haploids *SpC* that diploidized before mating and haploid *SpB* strains ^22^. WGD occurs also in L1 and L2 hybrids with lower frequency than in VL_C (3% and 2% respectively). These hybrids share the same *SpC* parents with VL_C crosses and the same *SpB* parents with VL_B crosses, suggesting that the WGD rate observed in L1 and L2 hybrids is potentially a trait inherited from the *SpC* parental sub-genome.

WGD occurs in hybrids but not in all biological replicates; only one of the two replicate crosses of the (M) and (H) crosses show WGD (M1 (4%) and H2 (6%)), suggesting that certain genotypes or combinations of genotypes are more prone to WGD (Fig. 1B, C). Because the parental strains used in M1 (*SpB1* and *SpA1*) and H2 (*SpB2* and *Scer2*) crosses are also involved in VL_B, VL_A and VL_S intra-lineage crosses in which WGD did not occur, our results cannot be explained by the ploidy instability of these parental strains, and thus support the hypothesis that WGD rate can be triggered by hybridization.

Genome doubling occurred at different generation timepoints after mating; some of them emerged quickly, in less than 90 generations, while others appeared after more than 680 generations (Fig. 1B, fig. S2). Two of the thirty-two tetraploids went extinct before the end of the experiment (L1_31 and H2_38). The remaining 30 tetraploids evolved from a few generations to more than 680 generations. We find that tetraploidy can be slowly reverted. One of the H2 tetraploids (H2_43) shows progressive reduction in ploidy from 4n to about 2.8n, 418 generations after WGD (fig. S2). Genome sequencing of this strain confirms several aneuploidies with copy number reductions for three large chromosomes (Chromosomes V, XII and XIV) (fig. S9 and S11). However, the copy number reduction of these three large chromosomes cannot fully explain such a decrease in ploidy (from 4n to 2.8n). These results could be explained by heterogeneity among isolated aneuploid colonies from glycerol stock, probably due to the high genomic instability of hybrids ^22^. Two of the M1 tetraploids (M1_32 and M1_40) also show rapid ploidy change from tetraploid to diploid. However, mixed colonies containing 4n and 2n strains in M1 lines have been detected and could explain this observation ^22^ (Fig. 1B, fig. S2).

Darlington (1937) ^12^ hypothesized that there should be an inverse relationship between the fertility of a diploid hybrid and that of a tetraploid to which it gives rise (Fig. 2A). He reasoned that at low parental divergence, a diploid cross would be fertile because homologous chromosomes will be able to pair at meiosis. However, the corresponding tetraploid would show low fertility because pairing could occur between any pair of the four homologous chromosomes, causing multivalent pairing and uneven segregation. At high parental divergence, the opposite would be expected. Diploid hybrids should be sterile due to the failure of homeologous chromosome pairing, but allopolyploids should be fertile due to consistent bivalent pairing between identical duplicated chromosomes at meiosis (Fig. 2 A).

**Fig. 2.**
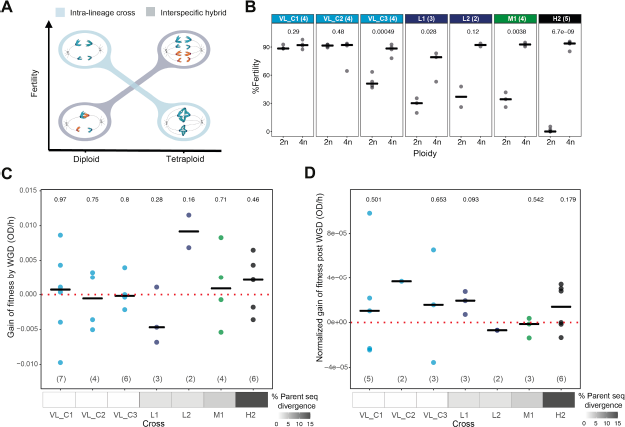
Whole genome duplication increases fertility and has no systematic trend towards higher fitness gain or loss in intra-lineage crosses and hybrids. (A) Darlington hypothesis: there is an inverse relationship between the fertility of a diploid hybrid and that of a tetraploid to which it gives rise. (B) Impact of whole genome duplication on fertility in intra-lineage crosses and hybrids. *P* values from a two-tailed paired t-test are shown. (C) WGD has no systematic trend towards immediate fitness gain or loss when it happens. Gain of fitness by WGD is calculated as the difference between the maximum growth rate before and after WGD. (D) Tetraploid lines show gains and losses of fitness at the end of the experiment. Gain of fitness post-WGD is calculated as the difference between the maximum growth rate after WGD and at the end of the experiment divided by the number of evolved generations following WGD to the end of the experiment. Medians are shown by horizontal bars. *P* values from a t-test are shown to evaluate if the gain of fitness is significantly different from zero. Numbers in parentheses represent sample sizes.

Previous studies from us and others confirmed that WGD can restore hybrid fertility ^20–22^. However, how WGD affects the fertility of intra-specific tetraploid crosses needs to be further tested. To test the first prediction of Darlington, we measured the fertility of 12 VL_C and all L (n = 5), M1 (n = 4) and H2 (n = 6) lines before and after WGD (the fertility of 8 of the L, M and H tetraploids were previously described ^22^). As expected, most allotetraploid hybrids have a significantly increased fertility compared to diploids that display low (L, M diploids) to almost null fertility (H diploids) (Fig. 2B). The only exception is observed for one tetraploid (H2_43) whose sterility is explained by a loss of sporulation ability. Contrary to what Darlington hypothesized and in agreement with previous studies in flowering plants ^29^, tetraploids from intra-specific crosses (VL_C1 and VL_C2 lines) show no difference between the diploid and tetraploid state in almost all tested lines, with both ploidy levels showing very high fertility (Fig. 2B). On the contrary, one VL_C3 line shows increased fertility after WGD, while VL_C lines that remain diploid show no change (fig. S5). These results suggest that there are accumulated genetic differences between the two parental *SpC* strains of the VL_C3 cross that decrease F1 fertility and which is restored by WGD. Indeed, these two parental genomes are more divergent compared to VL_C1 and VL_C2 (+2% and +4% SNPs respectively).

Several studies showed that polyploids are more successful than their diploid parents and undergo significantly faster adaptation, presumably because of their access to more mutations that lead to an increased phenotypic diversity ^7^. However, this increased rate of change could also increase the rate of deleterious mutations. To first test how WGD affects fitness under relaxed selection, we measured the maximum growth rate of the 32 lines that underwent WGD at 4 time points: soon after mating, before and after WGD, and at the end of the experiment (fig. S6). For comparison, we also measured the growth rate of 5 independent diploids from each cross after mating, in the middle and at the end of the experiment (fig. S6).

Our results show that spontaneous WGD leads to both gains and losses of fitness, such that WGD has no systematic trend towards higher fitness gain or loss in these conditions (Fig. 2C) (paired t-tests, (fig. S7A)). Evolved tetraploids in the majority of crosses also show gains and losses of fitness at the end of the experiment (Fig. 2D, fig. S7B). However, two of the H2 hybrids (H2_43 and H2_57) show respectively a markedly increased and decreased fitness (fig. S7B), suggesting that these hybrid lines had accumulated mutations that highly affected their growth rates and supporting the idea that allopolyploidy might increase phenotypic diversity.

It has been suggested that hybridization may increase the probability of WGD because combining two diverged genomes together would upset the course of cell division ^13^. Newly formed hybrids indeed have an increased rate of alterations in the genome (genomic instability, GIN) ^30,31^ and WGD is one of the common GIN hallmarks in cancer cells ^32,33^. However, little is known about the possible role of GIN in increasing the rate of WGD in hybrids. Our results show a higher rate of WGD in VL_C and L crosses, and only one of the two biological replicates of M (M1) and H (H2) hybrids. This result could be due to an overall increased GIN in crosses showing higher rates of WGD.

To test this hypothesis, we sequenced the genomes of n = 33-39 randomly chosen diploid lines from VL_B (1 and 2), VL_C, M and H crosses, as well as n = 34 diploid and n = 36 triploid lines from L crosses soon after mating and at the end of the experiment. We measured the rates of aneuploidy and loss of heterozygosity (LOH), two typical GIN hallmarks ^33,34^. We looked at the diploid lines of these crosses, reasoning that crosses with more intense GIN would also have more unstable diploid lines. Our results indicate that crosses showing higher rates of WGD manifest different hallmarks of increased GIN. For instance, diploid intra-lineage VL_C crosses showing the highest rates of WGD also exhibit the highest aneuploidy rate compared to VL_B crosses and to M and H hybrids (Kruskal Wallis and Mann-Whitney-Wilcoxon post-hoc pairwise comparisons, *P*= 1.4e-10, *P*= 7.7e-06 and *P*= 1.1e-07 respectively, Fig. 3A, B, fig. S8, S9). These crosses have at least one additional copy of chromosome XII inherited from their parental strains (fig. S10). The same result is observed for L hybrids, which share the same *SpC* parents with VL_C crosses (Kruskal Wallis and Mann-Whitney-Wilcoxon post-hoc pairwise comparisons, *P*= 0.0046 and *P*=0.039 respectively to VL_B crosses and H hybrids, Fig. 3A, B). This supports previous observations that the presence of extra chromosomes increases genomic instability ^35^ and could potentially trigger WGD by having additional chromosomes forming bridges that can prevent cytokinesis ^36^.

**Fig. 3.**
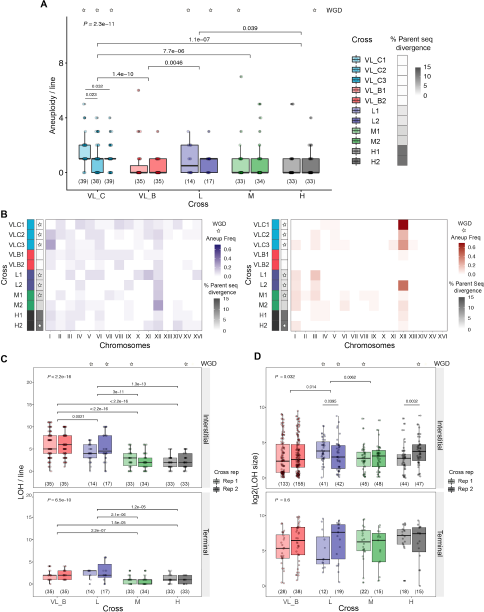
Higher WGD rate correlates with higher genomic instability. (A) Aneuploidy rate among diploid intra-lineage crosses and hybrids. (B) Different patterns in chromosome gain (left panel) and loss (right panel) in diploid intra-lineage crosses and hybrids. (C) Loss of heterozygosity (LOH) rate in diploid intra-lineage crosses and hybrids decreases with genetic divergence. (D) Interstitial and terminal LOH segments size in diploid intra-lineage crosses and hybrids. The crosses with a star are those where WGD occurred. Numbers in parentheses represent sample sizes. *P* values from Kruskal Wallis test (above) and pairwise Mann-Whitney-Wilcoxon tests are shown (only *P* values <0.05 are shown). For all boxplots the bold center line corresponds to the median value, the box boundaries correspond to the 25th and the 75th percentile, the whiskers correspond to 1.5 times the interquartile range.

Our results also show that all hybrids exhibit increased GIN compared to the intra-lineage crosses VL_B and VL_S2 (Table S2). Hybrids show statistically significant higher rates of whole chromosome loss (Fig. 3B, fig. S8) and a lower LOH frequency (Fig. 3C). This is consistent with a lower efficiency of mitotic homologous recombination between homeologous chromosomes which show a higher nucleotide divergence^37^. Aneuploidy frequencies of all 16 chromosomes indicate that chromosome XII is the most unstable chromosome among hybrids (Fig. 3B). This chromosome contains the rDNA locus, which is a large tandemly repeated sequence of 9kb encoding for ribosomal RNA, one of the most unstable parts of the genome due to the repeated recombinations and copy number variation ^38,39^. The high instability of this chromosome in hybrids could be a testimony of increased genomic instability.

Finally, we find that the replicates within crosses M and H that show higher rates of WGD (M1 and H2) manifest more hallmarks of GIN. The two replicates of interspecific hybrids, H1 and H2, show different LOH size ranges. The H2 hybrid lines (6% WGD) have significantly longer tract of interstitial LOH segments than H1 lines (0% WGD) (Mann-Whitney-Wilcoxon test, *P*= 0.0032, Fig. 3D). The lengths of these regions argue against a classical gene conversion mechanism that would in general lead to small LOH segments. Instead, they suggest a higher frequency of LOH caused by aberrant chromosomal segregation events and/or break-induced replication (BIR) events ^40^. Long-range LOH events have been described to be more frequent under replication stress and to have a greater contribution to GIN in aging and cancer cells ^41–43^. Finally, M1 (4% WGD) and M2 (0% WGD) hybrids show a significant difference in rates of line loss. Much higher line decline has been observed in M1 hybrids, suggesting that these hybrids have higher GIN leading to the frequent segregation of highly deleterious variants compared to M2 hybrids ^22^.

Increased GIN as a consequence of polyploidy was also previously observed ^2,7^ but not without the confounding effect of natural selection, which biases the set of visible mutations. We therefore investigated whether lines that went through WGD display higher genomic variability than diploids within the same cross. If these genomic changes occurred after WGDs this would suggest that WGDs actually increase the GIN. We find that frequencies of aneuploidy in tetraploids are up to four times those of diploids (Mann-Whitney-Wilcoxon test, *P*= 7.3e-05 Fig. 4A). Different patterns of chromosome gain and loss are observed in tetraploids compared to diploids (Fig. 3B, Fig. 4B, fig. S11, S12). Chromosome loss, consistent with previous studies (for review ^44^), more frequently affects the smallest chromosomes (I and III) in diploid hybrids, probably because it affects a smaller number of genes. However, larger chromosome losses (II, V and VIII) are observed in tetraploids, suggesting a potentially less deleterious effect of their loss due to the multiple copies present in the cell. Furthermore, these chromosome losses occurred most likely after WGD. Indeed, if a chromosome loss occurs before WGD then aneuploid lines would be homozygous for that chromosome. Allele frequency analysis of tetraploid lines show changes in allele frequency of aneuploid chromosomes but all of them remain heterozygous (fig. S13). These spontaneous aneuploidies have consequences on fitness. Two allopolyploids (H2_43 and H2_57) respectively show the most increased and decreased growth rates after WGD (fig. S7B) and exhibit multiple aneuploidies (3 and 4 aneuploid chromosomes while the average aneuploidy by line in this cross is about 2 aneuploidy) (fig. S11, S13). Tetraploids also show non-significant differences with diploids in LOH rate (Fig. 4C). However, they exhibit a higher number of the short-range LOH segments (Mann-Whitney-Wilcoxon test, *P*= 5.4e-04, Fig. 4D), indicating substantial differences in DNA repair process. It is worth noting that the LOH rate in tetraploids could be underestimated. In tetraploids, recombination events can only be detected between homeologous chromosomes, while recombination events between homologous chromosomes are undetectable. Consistent with this hypothesis, LOH analysis in triploid L1 and L2 hybrids show a decreased detectable LOH frequency leading to *SpC* parent allele loss compared to diploids (fig. S14). The presence of two identical copies of chromosomes in allopolyploids might increase the efficiency of homologous recombination, thus improving the DNA repair process during mitosis and consequently could contribute to genome stabilization. Overall, neo-polyploid hybrids are characterized by increased genomic variability that can affect fitness and suggest that WGD might be a springboard for hybrid genome stabilization in the long term.

**Fig. 4.**
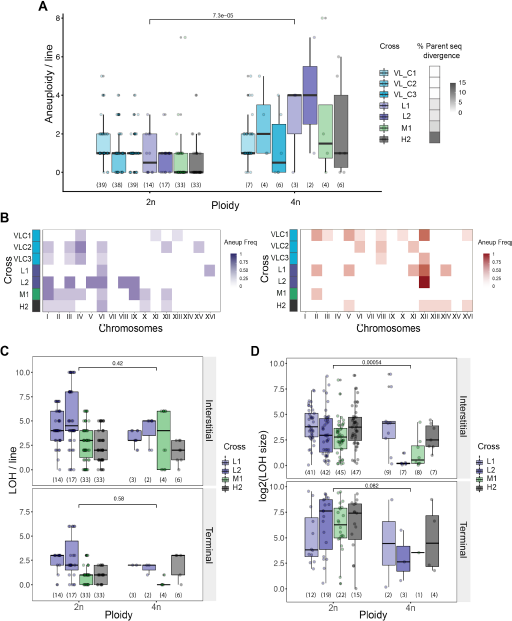
Tetraploid lines display increased genomic variability compared to diploids. (A) Aneuploidy frequency increases with ploidy in intra-lineage VL_C crosses and hybrids. (B) Patterns of chromosome gain (left panel) and loss (right panel) in tetraploid intra-lineage VLC crosses and hybrids. (C) Loss of heterozygosity (LOH) rate in tetraploids compared to diploid hybrids. (D) Interstitial and terminal LOH segment sizes in tetraploids compared to diploid hybrids. *P* values from Mann-Whitney-Wilcoxon test are shown. Numbers in parentheses represent sample sizes. For all boxplots the bold center line corresponds to the median value, the box boundaries correspond to the 25th and the 75th percentile, the whiskers correspond to 1.5 times the interquartile range.

Our results show that WGD occurs spontaneously in yeast hybrids within less than 90 cell divisions after mating, which may correspond to as few as 60 days in nature. We find that some lineages are more prone to WGD, and that WGD could be triggered by hybridization, most likely through increased genomic instability. GIN, whether intrinsic to some genotypes or triggered by hybridization, may accelerate WGD which in turns leads to increased fertility, genetic and phenotypic diversity that may contribute to the establishment of both autopolyploid and allopolyploid. There is therefore a positive feedback between hybridization and WGD in evolution, hybridization could increase the rate of WGD, which in turns leads to more instability that fuels further adaptation.

## Methods

### Experimental crosses

A total of 1304 lines from 15 different crosses were used; 864 of the lines from VL_B1/2, VL_C1/2/3, L1/2, M1/2, and H1/2 crosses and 152 of the lines from the VL_S2 cross were described previously ^22,26–28^. In this study, we generated 288 new lines (3 new crosses) VL_B3 (96 lines), VL_A (96 lines) and VL_S1 (96 lines). These were generated as previously described by Charron et al., (2019)^22^. Haploids were precultured overnight in 5 mL of YPD (1% yeast extract, 2% tryptone and 2% D-glucose). All incubation steps were performed at room temperature. Precultures were then diluted at OD600nm of 1.0 in 500 μL aliquots. The aliquots from two strains to be crossed were mixed together and 5 μL were used to inoculate 200 μL of fresh YPD medium in 96 replicates so all individual strains derived from independent mating events and would be truly independent hybrids. Cells were given 6 hours to mate after which 5μL of the mating cultures were spotted on a diploid selection medium (YPD, 100 μg mL-1 G418, 10 μg mL-1 Nourseothricin). From each of the 96 spots, one colony was picked as a founding line for the evolution experiment, resulting in 96 independent lines for VL_B3, VL_A and VL_S1.

### Evolution experiment

The 288 lines generated were evolved as previously described by Charron et al., (2019)^22^. Each of the independent lines (single colonies) were streaked on a sector corresponding to one-third of a YPD agar plate. Plates were incubated at room temperature for 3 days after which a new single colony was streaked as a progenitor for the new generation. Every 3 passages, the colonies were both streaked and used to inoculate the wells of a 96 wells plate containing 150 μL of fresh YPD medium. After a 24 h of incubation at room temperature, 75 μL of 80% glycerol was added and the plates were placed in a −80 °C freezer for archiving. The lines were maintained on plates for a total of 35 passages, which is about ~770 generations ^22^.

### Determination of ploidy

Measurement of the cell DNA content was performed using flow cytometry with the SYTOX™ green staining assay (Thermo Fisher, Waltham, USA) as done in (Charron et al., 2019)^22^. Cells were first thawed from glycerol stocks on solid YPD in omnitray plates (room temperature, 3 days) including controls. The parental strains *S. paradoxus SpB* strain (MSH604) and the *S. cerevisiae* strain (LL13_054) were used as controls on both their haploid and diploid (wild strain) state. Liquid YPD cultures of 1?ml in 96-deep-well (2 ml) plates were inoculated and incubated for 24?h at room temperature. Cells were subsequently prepared as in Gerstein et al. (2006)^45^. Cells were first fixed in 70% ethanol for at least 1?h at room temperature. RNAs were eliminated from fixed cells using 0.25?mg/ml of RNAse A during an overnight incubation at 37?°C. Cells were subsequently washed twice using sodium citrate (50 mM, pH7) and stained with a final SYTOX™ green concentration of 0.6 μM for a minimum of 1?h at room temperature in the dark. The volume of cells was adjusted to be around a cell concentration of less than 500?cells?μL−1. Five thousand cells of each sample were analyzed on a Guava^^®^^ easyCyte 8HT flow cytometer using a sample tray for 96-well microplates. Cells were excited with the blue laser at 488?nm and fluorescence was collected with a green fluorescence detection channel (peak at 512?nm). The distributions of the green fluorescence values were processed to find the two main density peaks, which correspond to the two cell populations, respectively in G1 and G2 phases. The data were analysed using R version 3.6.1. The code is available at https://github.com/Landrylab/Marsit_et_al_2020.

### Measurement of fertility

Strains were thawed and 2 μL of the stocks were spotted on a fresh YPD medium and incubated for 3 days. A small number of cells were used to inoculate 4 mL of fresh YPD media and incubated for another day. From those precultures, a new 4 mL culture was inoculated at OD595 of 0.6 in fresh YPD and grown for 3 hours. Cells were subsequently prepared as in Charron et al. (2019)^22^. Cell cultures were centrifuged, the YPD was replaced with 4 mL of YEPA medium (1% yeast extract, 2% tryptone and 2% potassium acetate). Cultures were incubated for 24 h after which they were centrifuged again, washed once with sterile deionized water, and put into 4 mL of SP medium (0.3% potassium acetate 0.02% d-Raffinose). After 5-7 days of incubation, the strains were dissected as in Charron et al. (2019)^22^ with a SporePlay™ dissection microscope (Singer Instruments, Somerset, UK) on YPD plates and incubated for 5 days at 25 °C. Pictures of the plates were taken after the incubation time and fertility was determined as the number of spores forming a colony visible to the naked eye after 5 days.

### Growth rate measurement

A total of 233 strains (32 tetraploids at 4 time points and 35 diploids randomly selected at 3 timepoints) were thawed from glycerol stocks on solid YPD omnitray plates (25 °C, 72 h). Four to five independent replicates from each strain were pre-cultured in 1 mL of YPD liquid cultures in 96 deep-well plates (2 ml) and incubated for 24 h at 25 °C. Precultures were then diluted to OD595 = 0.1 and incubated at 25 °C to reach the OD595 = 0.6. Subsequently, 20 μL of these pre-cultures were grown in 96-well flat-bottomed culture plates in 180 μL of media (YPD), resulting in an initial OD595 of approximately 0.1. Incubation at 25 °C was performed directly in 3 temperature-controlled spectrophotometers (Infinite^®^ 200 PRO, Tecan, Reading, UK) that read the OD595 at intervals of 15 min. The growth rate of each replicate was estimated from growth curves using R v3.6.1 (the code is available at https://github.com/Landrylab/Marsit_et_al_2020). The growth rate was computed as the 98th percentile of the set of linear regression slopes fitted in ten-timepoint wide overlapping sliding windows.

### Whole-genome sequencing

We performed whole-genome sequencing of 864 strains:

1. 33 to 40 randomly selected diploid lines from VL_B1, VL_B2, VL_C1, VL_C2, VL_C3, M1, M2, H1, H2;
2. 13 and 16 diploid and 19 and 17 triploid lines from L1 and L2;
3. all of the 32 strains that because tetraploid were sequenced soon after mating and at the end of the experiment (and at the middle of the experiment after 352 generations for tetraploids that show ploidy change to diploid);
4. the 13 corresponding haploid parental strains.

Genomic DNA was extracted from overnight cultures from one isolated colony of each stock following standard protocols (QIAGEN DNAeasy, Hilden, Germany). Extracted DNA was treated with RNase A and purified on Axygen AxyPrep Mag PCR Clean-up SPRI beads. Libraries were prepared with the Illumina Nextera kit (Illumina, San Diego, USA) following the manufacturer's protocol and modifications from ^46^. The quality of a few randomly selected libraries was controlled using an Agilent BioAnalyzer 2100 electrophoresis system. Pooled libraries were sequenced in paired-end, 150 bp mode on different lanes of HiSeqX (Illumina, San Diego, USA) at the Genome Quebec Innovation Center (Montréal, Canada). The 864 genomes were sequenced with an average genome-wide coverage of 90X. Raw sequences are accessible at NCBI (bio project ID PRAJNA 515073).

### Read mapping

Raw reads were trimmed using Trimmomatic version 0.33 with parameters ILLUMINACLIP:nextera.fa:6:20:10 MINLEN:40 and a library of Illumina Nextera adapter sequences. Reads were subsequently mapped on the reference genome of *S. paradoxus* MSH604 strain, one of the four used SpB parental strains, for VL_B crosses and all hybrids, onto the *SpC* parental strain (LL2011_012) for the VL_C crosses, and onto *S. cerevisiae* YPS128 strain for H hybrids using bwa mem v0.7.17. Mapped reads were sorted using samtools sort version 1.8. and duplicates were removed using Picard RemoveDuplicates version 2.18.29-SNAPSHOT with parameter REMOVE_DUPLICATES=true.

### Read depth

Read coverage for each position in the genome was estimated using SAMtools depth v1.8. and averaged over 10kb windows to detect copy number variation within and among chromosomes. The read depth of coverage obtained for each bin of 10 kb on each chromosome was divided by the whole genome coverage. The median of the values obtained for each chromosome corresponds to the chromosome copy number and the difference between these values at T_end_ (at the end of the experiment) and T_ini_ (soon after mating) corresponds to the number of gained or lost chromosomes. The data was analyzed using R version 3.6.1 (the code is available at https://github.com/Landrylab/Marsit_et_al_2020).

### Allele frequency and LOH

Single nucleotide polymorphisms (SNP) were called using Freebayes v1.3.1 ^47^ accounting for different ploidy levels of strains and chromosomes. Control-FREEC v11.5 ^48,49^ was used to determine copy number in 250 bp non-overlapping windows of each genome with options breakPointThreshold = 0.8, minExpectedGC = 0.35, maxExpectedGC = 0.55, telocentromeric = 7000. The most common copy number occurring across windows for a given chromosome in a given strain was set as the copy number of that strain and that chromosome. Variant calling was run for each cross replicate separately including both haploid parental strains. The following options were used: -q 20 --use-best-n-alleles 4 --limit-coverage 20000 -F 0.02. Multi-nucleotide polymorphisms were decomposed into SNPs using script vcfallelicprimitives with -kg option. The criteria for selecting SNPs included QUAL > 1, QUAL / AO > 10 (where QUAL is a quality of variant and AO is the number of alternate alleles), SAF > 0 (number of alternate observations on the forward strand), SAR > 0 (number of alternate observations on the reverse strand), RPR > 1 (number of reads centered to the right of an alternate allele), RPL > 1 (number of reads centered to the left of an alternate allele), MQM / MQMR > 0.9 and MQM / MQMR < 1.05 (where MQM and MQMR are mean mapping qualities of alternate and reference alleles, respectively). Indels, multiallelic SNPs, and SNPs overlapping repeats were excluded.

For allele frequency calculation, only heterozygous loci between parental genomes for each cross were kept and the parental origin of each allele was identified. Allele frequencies of heterozygous loci were calculated as the allele read depth corresponding to one of the two parents divided by the total read depth of the locus. Only loci with read depth higher than 20 reads were considered. Loss of heterozygosity tracts were identified. SNPs with allele frequencies deviating from the average allele frequency over the whole chromosome (+/− 0.15) were kept as potential SNPs belonging to an LOH tract. Stretches of consecutive marker positions were grouped in LOH regions. Blocks of LOH were identified by looking for a minimum of 3 successive SNPs with the same allele frequency (+/− 0.1) in a window of 300 bp for hybrids and a window of 1 kb for VL_B crosses. The size of LOH segments was calculated as the difference between the position of the last and the first SNP of each identified LOH segment. Because the minimum LOH size was of 1 kb in VL_B crosses due to the very low heterozygosity in these crosses, we considered in our comparisons of LOH frequencies between crosses only LOH segments larger than 1 kb. The data was analyzed using R version 3.6.1 (the code is available at https://github.com/Landrylab/Marsit_et_al_2020).

### Statistical analyses

Statistical analyses and figure creation for all data were done using custom scripts available at https://github.com/Landrylab/Marsit_et_al_2020 in R version 3.6.1.

## Acknowledgments

We thank H. Martin for contributing to preliminary sequence data analyses. We also thank the members of the Landry lab for discussions and M. Barker, A.M. Dion-côté, C. Bautista Rodriguez, A. F. Cisneros and D. Ascencio, for comments on the manuscript.

## Author contributions

Conceptualization, C.R.L. and S.M.; Data curation, S.M. and M.H; Funding acquisition, C.R.L.; Experimental work: S.M., M.H. and G.C., Formal analysis: S.M. and A.F. Project supervision, C.R.L.; Writing - original draft, S.M and C.R.L..; Writing – review & editing, all authors.

## Data and materials availability

Raw sequencing data that support the findings of this study have been deposited in Sequence Read Archive (SRA), NCBI with the BioProject accession code #PRJNA515073. Flow cytometry data that support the findings of this study are available in figshare with the identifier Marsit_et_al_Ploidy_data_2020. All yeast strains are available from C.R.L. under a material transfer agreement.

## Supplementary Materials

**Fig. S1.**
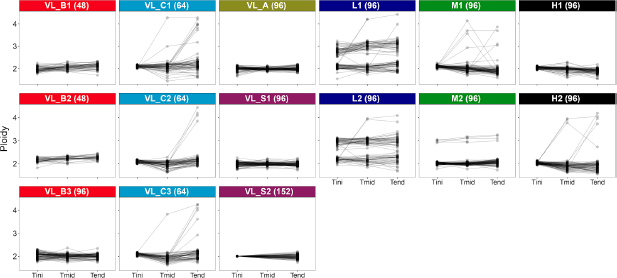
Ploidy evolution of the 1304 lines from the 15 different crosses at three different generation time points. T_ini_ (22 generations), T_mid_ (352 generations) and T_end_ (770 generations and 2062 generations for VL_S2). The numbers in parentheses represent sample sizes.

**Fig. S2.**
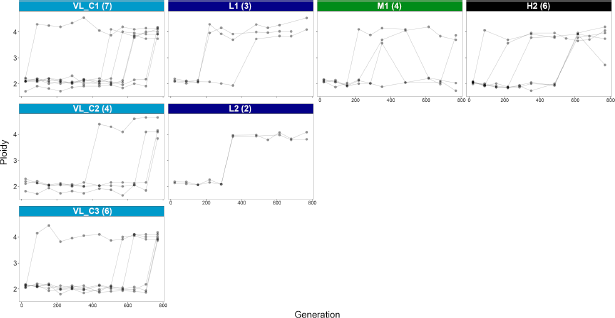
Ploidy evolution each ~70 generations of the 32 lines where WGD occurred shows that WGD occurred at different generation time points. The numbers in parentheses represent sample sizes.

**Fig. S3.**
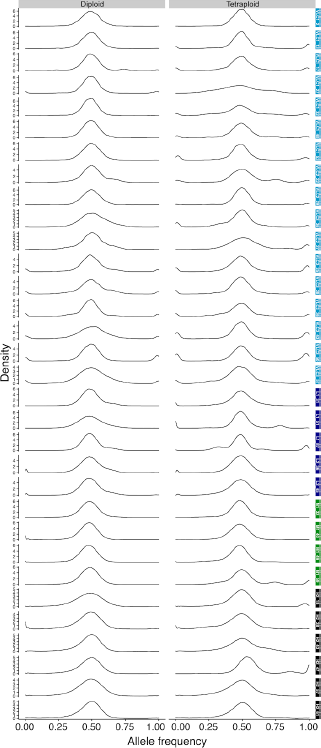
Allele frequency over the 16 chromosomes of the 32 identified lines that went through WGD at their diploid and tetraploid state confirm that both parental genomes have been duplicated.

**Fig. S4.**
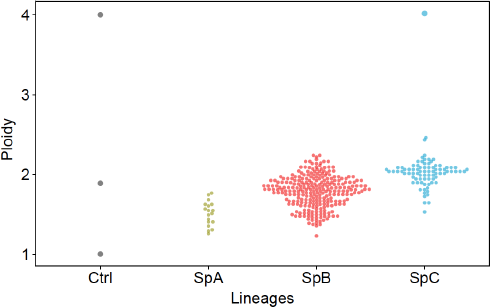
A tetraploid *SpC* natural isolate is identified among natural *S. paradoxus* strains. The ploidy of 366 wild Noth-American *S. paradoxus* isolates from *SpA* (n= 18), *SpB* (n= 265) and *SpC* (n= 83) lineages were estimated using flow cytometry. Ctrl corresponds to MSH604 *SpB* parent at haploid and diploid (wild) states used as controls.

**Fig. S5.**
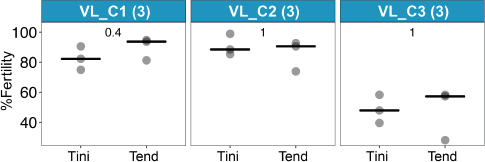
The fertility of F1 hybrids from intra-lineage crosses (VL_C diploids) following mating and at the end of the experiment shows that there is no systematic trend towards fertility gain or loss. *P*-values from a two-tailed paired t-test are shown. The numbers in parentheses represent sample sizes.

**Fig. S6.**
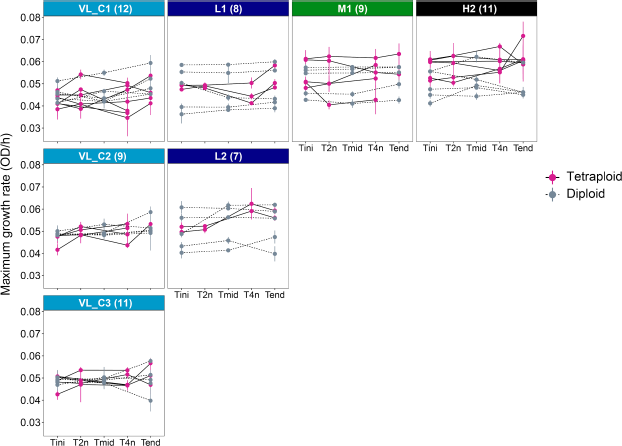
WGD has no systematic trend toward fitness gain or loss. The maximum growth rate of 32 tetraploids at 4 time points and 35 diploids randomly selected at 3 time points were evaluated. The maximum growth rate of tetraploids was evaluated following mating (T_ini_), before WGD (T_2n_), after WGD (T_4n_), and at the end of the experiment after 770 generations (T_end_). Maximum growth rate of diploids was evaluated following mating (T_ini_), at the middle of the experiment after 352 generations (T_mid_) and at the end of the experiment after 770 generations (T_end_). Four to five independent replicates have been performed for each line. Error bars represent SD. The numbers in parentheses represent sample sizes.

**Fig. S7.**
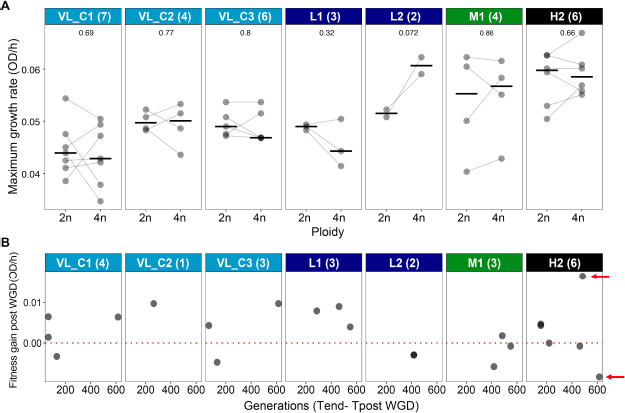
Tetraploid lines show gains and losses of fitness following WGD and at the end of the experiment. (A) Maximum growth rate before (2n) and after WGD (4n) of intra-lineage VL_C crosses and hybrids. *P* values from a two-tailed paired t-test are shown above. Medians are shown by horizontal bars (B) Fitness gain of tetraploids at the end of the experiment. Fitness gain post WGD is calculated as the difference of the maximum growth rate between the end of the experiment and following WGD. Evolved generations are calculated as the difference between the number of generations following WGD (Tpost WGD) and the end of the experiment (T_end_). Red arrows indicate the H2_43 and H2_57 lines, showing respectively, markedly increased and decreased fitness. The numbers in parentheses represent sample sizes.

**Fig. S8.**
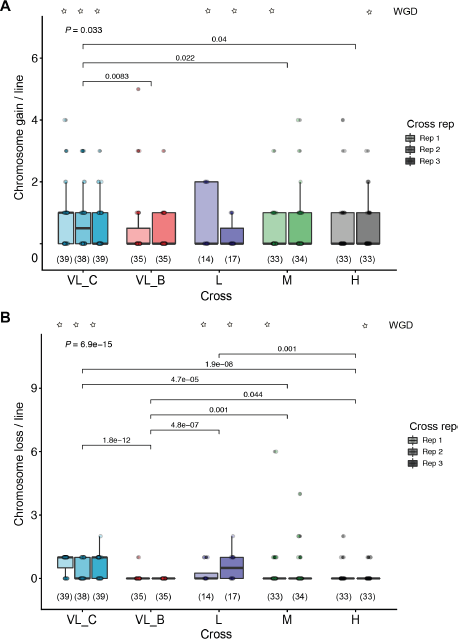
Chromosome gain and loss in intra-lineage crosses and hybrids. (A) Chromosome gain rate in intra-lineage crosses and hybrids. (B) Chromosome loss rate in intra-lineage crosses and hybrids. *P* values from Kruskal Wallis test (above) and pairwise Mann–Whitney–Wilcoxon tests are shown (only *P* values <0.05 are shown). The crosses with a star are those where WGD occurred. The numbers in parentheses represent sample sizes. For all boxplots the bold center line corresponds to the median value, the box boundaries correspond to the 25th and the 75th percentile, the whiskers correspond to 1.5 times the inter-quartile range.

**Fig. S9.**
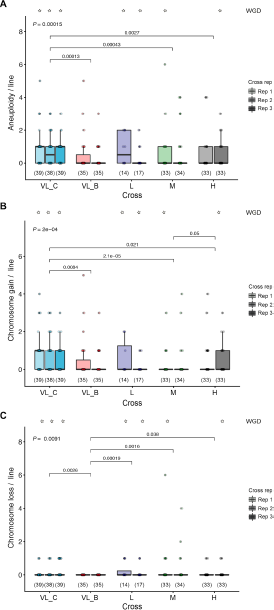
Chromosome gain and loss in intra-lineage crosses and hybrids including all chromosomes except chromosome 12. (A) Aneuploidy rate excluding chromosome 12 in intra-lineage crosses and hybrids. (B) Chromosome gain rate excluding chromosome 12 in intra-lineage crosses and hybrids. (C) Chromosome loss rate excluding chromosome 12 in intra-lineage crosses and hybrids. Numbers in parentheses represent sample sizes. *P* values from Kruskal Wallis test (above) and pairwise Mann–Whitney–Wilcoxon tests are shown (only *P* values <0.05 are shown). The crosses with a star are those where WGD occurred. The numbers in parentheses represent sample sizes. For all boxplots the bold center line corresponds to the median value, the box boundaries correspond to the 25th and the 75th percentile, the whiskers correspond to 1.5 times the inter-quartile range.

**Fig. S10.**
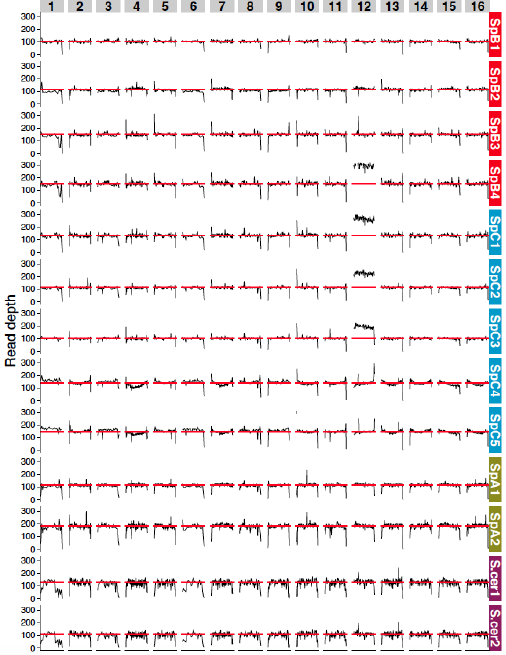
Read depth over the 16 chromosomes of parental strains used for intra-lineage and inter-specific crosses show an additional copy of chromosome 12 in three *SpC* and one *SpB* parents. Black lines correspond to the read depth over windows of 10kb across chromosomes and the red lines correspond to the coverage of the whole genome.

**Fig. S11.**
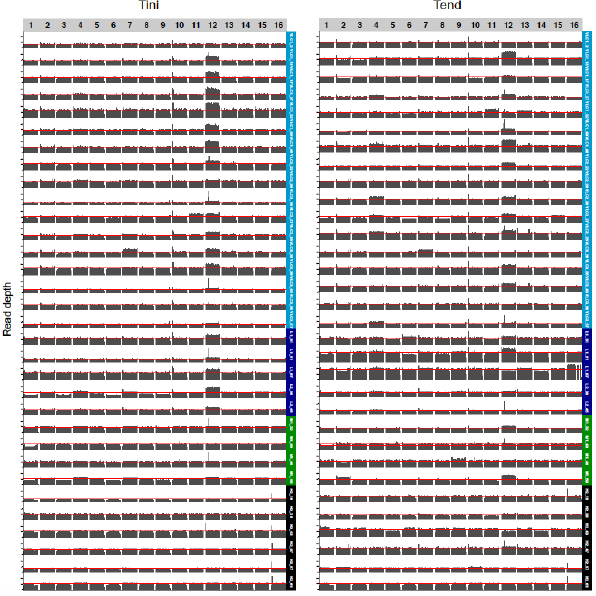
Read depth over the 16 chromosomes of the 32 lines where WGD occurred following mating (T_ini_, diploids), and at the end of the experiment (T_end_, tetraploids) reveal several aneuploidies. Black bars correspond to the read depth over windows of 10 kb across chromosomes and the red lines correspond to the coverage of the whole genome.

**Fig. S12.**
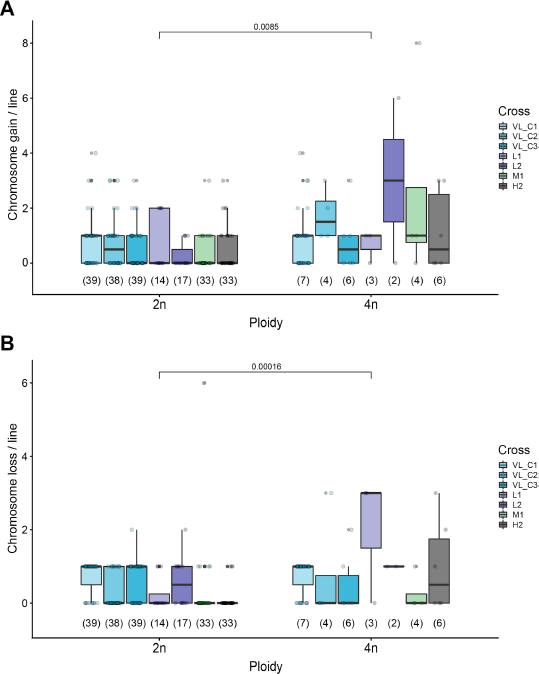
Chromosome gain and loss rate comparison between diploid and tetraploid intra-lineage VL_C crosses and hybrids. (A) Chromosome gain rate comparison between diploid and tetraploid intra-lineage VL_C crosses and hybrids. (B) Chromosome loss rate comparison between diploid and tetraploid intra-lineage VL_C crosses and hybrids. *P* values from Mann–Whitney–Wilcoxon test are shown above. The numbers in parentheses represent sample sizes. For all boxplots the bold center line corresponds to the median value, the box boundaries correspond to the 25th and the 75th percentile, the whiskers correspond to 1.5 times the inter-quartile range.

**Fig. S13.**
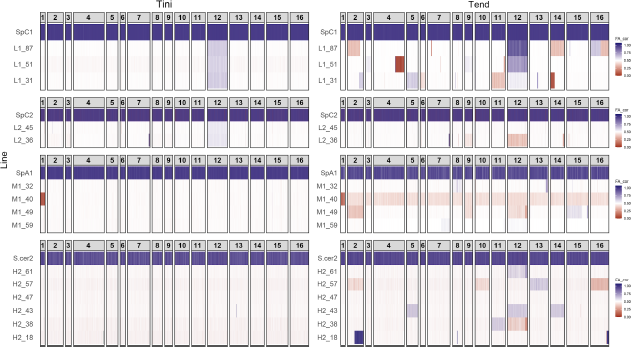
Allele frequency over the 16 chromosomes in hybrid tetraploids reveals many LOH events and variation in parental contributing allele frequencies resulting from aneuploidies following mating (T_ini_) and at the end of the experiment after 770 generations (T_end_).

**Fig. S14.**
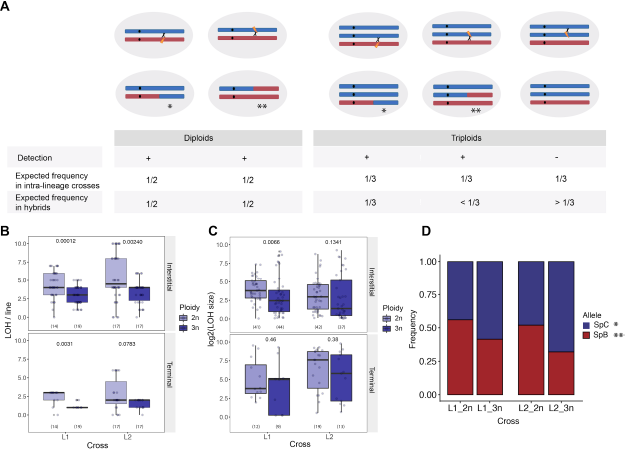
The presence of two identical copies of chromosomes in allopolyploids leads to a decrease in mitotic recombination between homeologous chromosomes (A) The different LOH possibilities and their expected frequency in triploids compared to diploids. (B)The frequency of detectable LOH in triploid L1 and L2 hybrids is lower than in diploids. (C) LOH segments size in triploids compared to diploids L1 and L2 hybrids. (D) LOH leading to segments with homozygous *SpC* alleles are more frequent than those leading to an increase in *SpB* allele frequency. LOH events leading to segments with *SpC* alleles are labeled with one star (*) and LOH events leading to segments with *SpB* alleles are labeled with two stars (**). Numbers in parentheses represent sample sizes. *P* values from Mann–Whitney–Wilcoxon test are shown above. For all boxplots the bold center line corresponds to the median value, the box boundaries correspond to the 25th and the 75th percentile, the whiskers correspond to 1.5 times the inter-quartile range.

**Table S1.** List of crosses used in this study

| **Cross** | **a strain** | **a strain**<br>**resistance** | **$\alpha$ strain** | **$\alpha$ strain**<br>**resistance** | **Total number** | **Reference** |
| --- | --- | --- | --- | --- | --- | --- |
| VL_B1 | MSH604 | NAT | LL12_028 | G418 | 48 | Charron et al.,<br>2019 |
| VL_B2 | UWOPS_91_202 | G418 | LL12_021 | NAT | 48 | Charron et al.,<br>2019 |
| VL_B3 | MSH604 | NAT | UWOPS_91_<br>202 | G418 | 96 | This study |
| VL_C1 | LL11_004 | NAT | MSH587 | G418 | 64 | Hénault et al., 2020 |
| VL_C2 | LL11_009 | G418 | LL11_001 | NAT | 64 | Hénault et al., 2020 |
| VL_C3 | LL11_009 | NAT | LL11_012 | G418 | 64 | Hénault et al., 2020 |
| VL_A | YPS744 | NAT | YPS644 | G418 | 96 | This study |
| VL_S1 | LL13_040 | G418 | LL13_054 | NAT | 96 | This study |
| VL_S2 | BY4741 | - | BY4742 | - | 152 | Hall et al., 2008 |
| L1 | MSH604 | NAT | LL11_004 | G418 | 96 | Charron et al.,<br>2019 |
| L2 | UWOPS_91_202 | G418 | LL11_009 | NAT | 96 | Charron et al.,<br>2019 |
| M1 | MSH604 | NAT | YPS644 | G418 | 96 | Charron et al.,<br>2019 |
| M2 | UWOPS_91_202 | G418 | YPS744 | NAT | 96 | Charron et al.,<br>2019 |
| H1 | MSH604 | NAT | LL13_040 | G418 | 96 | Charron et al.,<br>2019 |
| H2 | UWOPS_91_202 | G418 | LL13_054 | NAT | 96 | Charron et al.,<br>2019 |

**Table S2.** Rates of WGD, aneuploidy, and LOH by generations of intra-lineage crosses and hybrids.

| Cross | Ploidy | WGD rate /gen | Aneuploidy rate /gen | Chromosome gain rate /gen | Chromosome loss rate /gen | LOH rate /gen |
| --- | --- | --- | --- | --- | --- | --- |
| VL_B1 | 2n | 0 | 6.88 +/- 16.3 E-04 | 6.49 +/- 14.6 E-04 | 0.38 +/- 2.26 E-04 | 7.23 +/- 4.09 E-03 |
| VL_B2 | 2n | 0 | 4.97 +/- 8.63 E-04 | 4.97 +/- 8.63 E-04 | 0 | 6.95 +/- 3.56 E-03 |
| VL_B3 | 2n | 0 | - | - | - | - |
| VL_A | 2n | 0 | - | - | - | - |
| VL_S1 | 2n | 0 | - | - | - | - |
| VL_S2 | 2n | 0 | 1.04 +/- 1.84 E-04 * | 0.97 +/- 0.18 E-04 * | 0.7 +/- 0.04 E-05 * | - |
| VL_C1 | 2n | 1.42E-04 | 2.02 +/- 1.4 E-03 | 10.3 +/- 1.28 E-04 | 9.94 +/- 5.91 E-04 | - |
| VL_C2 | 2n | 8.11E-05 | 1.41 +/- 1.4 E-03 | 10.2 +/- 1.3 E-04 | 3.87 +/- 6.14 E-04 | - |
| VL_C3 | 2n | 1.21E-04 | 1.44 +/- 1.2 E-03 | 7.2 +/- 12.8 E-04 | 7.2 +/- 7.4 E-04 | - |
| L1 | 2n | 4.05E-05 | 1.23 +/- 1.4 E-03 | 8.91 +/- 13.2 E-04 | 3.34 +/- 6 E-04 | 5.47 +/- 2.26 E-03 |
| L2 | 2n | 2.70E-05 | 1.07 +/- 1.04 E-03 | 3.57 +/- 6.12 E-04 | 7.64 +/- 8.6 E-04 | 6.95 +/- 3.87 E-03 |
| M1 | 2n | 5.41E-05 | 8.91 +/- 1.7 E-03 | 4.46 +/- 8.63 E-04 | 4.46 +/- 14.4 E-04 | 3.61 +/- 2.16 E-03 |
| M2 | 2n | 0 | 1.14 +/- 1.8 E-03 | 7.08 +/- 12.4 E-04 | 4.33 +/- 11.3 E-04 | 3.33 +/- 2.01 E-03 |
| H1 | 2n | 0 | 7.7 +/- 16.4 E-04 | 6.08 +/- 12.1 E-04 | 1.62 +/- 5.55 E-04 | 2.8 +/- 1.56 E-03 |
| H2 | 2n | 8.11E-05 | 7.7 +/- 12.5 E-04 | 6.48 +/- 10.6 E-04 | 1.22 +/- 3.9 E-04 | 3.23 +/- 1.9 E-03 |
| VL_C1 | 4n | - | 2.02 +/- 1.4 E-03 | 1.03 +/- 1.28 E-03 | 9.94 +/- 5.91 E-04 | - |
| VL_C2 | 4n | - | 3.34 +/- 2.56 E-03 | 2.34 +/- 8.63 E-03 | 10 +/- 20 E-04 | - |
| VL_C3 | 4n | - | 1.78 +/- 2.3 E-03 | 1.11 +/- 1.56 E-03 | 6.68 +/- 11.2 E-04 | - |
| L1 | 3n | - | 1.8 +/- 1.2 E-03 | 0.56 +/- 6.78 E-03 | 1.27 +/- 1.1 E-03 | 3.56 +/- 1.78 E-03 |
| L1 | **4n** | - | 3.57 +/- 3 E-03 | 0.89 +/- 7.72 E-03 | 2.67 +/- 2.3 E-03 | 4.01 +/- 1.2 E-03 |
| L2 | **3n** | - | 2.91 +/- 3.2 E-03 | 1.67 +/- 2.21 E-03 | 1.34 +/- 1.5 E-03 | 4.06 +/- 2.11 E-03 |
| L2 | **4n** | - | 5.35 +/- 5.6 E-03 | 4.01 +/- 5.67 E-03 | 1.34 +/- 0 E-03 | 4.55 +/- 2.29 E-03 |
| M1 | **4n** | - | 3.68 +/- 4.8 E-03 | 3.34 +/- 4.9 E-03 | 3.34 +/- 6.6 E-04 | 6.09 +/- 2.92 E-03 |
| H2 | **4n** | - | 2.90 +/- 3.5 E-03 | 1.56 +/- 1.97 E-03 | 1.34 +/- 1.6 E-03 | 3.04 +/- 1.21 E-03 |

\* (Zhu et al., 2014) (50)

**Data S1.**

Zip file containing data for fertility and OD data for cell growth used in this study

